# The Slowest Timescales of Neural Synchronization Reveal the Strongest Influence of Auditory Distraction

**DOI:** 10.1101/2025.05.05.652235

**Authors:** David O. Sorensen, Jenna A. Sugai, Aravindakshan Parthsarathy, Kenneth E. Hancock, Daniel B. Polley

## Abstract

Among all the sounds occurring at any given time, people are often interested in listening to just one. Some competing sounds are merely background noise, whereas others distract attention from target sounds and are less easily suppressed. During active listening, the central auditory pathway unmixes target and distractor sounds based on temporal differences that vary across three orders of magnitude – from millisecond differences in acoustic temporal fine structure to slower perceptual grouping factors that stretch out to multiple seconds. Here, we developed an approach to directly measure central auditory encoding of multiplexed target and distractor sound features in human listeners to determine which timescales are most impacted by the presence of distracting sounds. Target sounds contained nested features along four timescales, including temporal fine structure (∼500 Hz), temporal envelope (∼25-80 Hz), envelope changes (∼5 Hz), and slower changes in embedded context reflecting whether target stimuli were randomly arranged or formed a repeating pattern (∼0.5 Hz). Targets were presented with competing sounds that provided variable levels of distraction: either a highly distracting melody or a less distracting noise. Neural synchronization to each timescale was simultaneously and independently measured for target and distractor sounds from electroencephalogram (EEG) recordings during a listening task. Sustained shifts from random to regular arrangements of temporal sequences were reliably perceived, yet did not evoke a pattern recognition potential, nor neural synchronization changes at any timescale. Synchronization to relatively slow changes in envelope transitions (<10Hz) of the target sound deteriorated with the addition of a more distracting sound while synchronization to more rapid fluctuations in the fine structure or envelope modulation rate were unaffected by varying levels of distraction. Categorizing trials according to task performance revealed a conjunction of enhanced entrainment to slower temporal features in the distractor sound and reduced synchronization to the target sound on error trials. By designing a stimulus paradigm that leveraged the remarkable temporal processing capabilities of the auditory nervous system, we were able to simultaneously quantify multiple target and distractor sound features reproduced in the EEG. This paradigm identified synchronization processes in the 7-10 Hz alpha range that has been linked to distractor suppression, which may prove valuable for research on clinical populations who report difficulty suppressing awareness of distracting sounds.

## 1 Introduction

Listening requires people to suppress all the sounds they are hearing except for the sound source in their attentional spotlight. Often characterized in the realm of speech-on-speech masking as the “cocktail party problem,” the simultaneous presentation of multiple auditory objects challenges the abilities of the auditory system together with central mechanisms of attention in order to selectively attend to an intended object and inhibit task-irrelevant objects (Cooke, Garcia Lecumberri and Barker, 2008). Difficulties in this domain may contribute to difficulties understanding speech in noisy environments, a common complaint of individuals claiming a hearing difficulty without hearing threshold shifts (Tremblay *et al*., 2015; Parthasarathy *et al*., 2020; Cancel *et al*., 2023).

To successfully attend to one sound in an auditory scene, the incoming auditory signal must be separated to form distinct auditory objects. Temporal cues that group together across different timescales are key features utilized by the auditory nervous system to separate different auditory objects (Sollini *et al*., 2022). More distracting background environments may degrade the encoding of intended target temporal features (Choi *et al*., 2014) such that measurements of temporal encoding across timescales in distracting environments would provide insight into auditory distraction.

The high temporal fidelity of the auditory nervous system allows measurements of synchronization to stimulus features (Galambos, Makeig and Talmachoff, 1981). Measurements of this synchronization, commonly called following responses, have been done in response to stimuli ranging from simple tones to repeated syllables. These include synchronization to the temporal fine structure of stimuli, termed frequency following responses (FFRs; see Krizman and Kraus, 2019 for a review), and synchronization to the amplitude envelope of stimuli, termed envelope following responses (EFRs; see Shinn-Cunningham et al., 2017 for review). Most frequently, studies of these following responses measure and report on responses to a single feature of the stimuli, but natural stimuli often contain multiplexed temporal features (Rosen, 1992). Studies of following responses also typically present the stimuli in quiet or acoustically simple background noise (Zhu *et al*., 2013; Bharadwaj *et al*., 2015), thus providing no insight into how temporal synchronization may be challenged by difficult listening environments.

While speech is the canonical stimulus for listening in a noisy environment, the variability in speech stimuli, especially naturally produced speech, reduces the ability to measure synchronization to the various features with specificity. Following responses require hundreds of trials to average to reliably measure responses (Bharadwaj and Shinn-Cunningham, 2014). Isolated syllables can be presented repeatedly to measure an FFR (Kraus *et al*., 2016), but the fundamental frequency of the temporal fine structure in natural speech varies across a range of hundreds of hertz (Stevens, 1971) precluding averaging.

In contrast, simple stimuli can be designed with well-controlled properties that can be repeated across presentations. This allows for synchronization to be measured even to stimulus features beyond temporal fine structure and amplitude envelopes, such as frequency modulation (Parthasarathy *et al*., 2020). Behaviorally, simple stimuli have often been presented under passive listening conditions (Zhu *et al*., 2013; Bharadwaj and Shinn-Cunningham, 2014; Barascud *et al*., 2016; Herrmann and Johnsrude, 2018; Herrmann, Buckland and Johnsrude, 2019; Calcus, Undurraga and Vickers, 2022) or in detection-based tasks (Durlach, Mason, Shinn-Cunningham, *et al*., 2003) but more complex perceptual judgments can be utilized as well (Southwell *et al*., 2017). Competitors can also be designed to minimize energetic masking in the auditory periphery (Durlach et al., 2003)—an important contributor to difficulty hearing in noisy environments but which is confounded with cognitive effects such as distraction (Brungart *et al*., 2001; Kidd *et al*., 2016).

Using novel stimuli, we report here a paradigm to simultaneously measure synchronization across multiplexed timescales in various levels of distraction. This paradigm allows us to probe the neurophysiological effects of auditory distraction on target encoding and measure distraction- sensitive features which may prove useful as objective markers of susceptibility to auditory distraction.

## 2 Materials and Methods

### 2.1 Participants

Participants were recruited from the general population via word of mouth, flyers, and advertisements on the Mass General Brigham participant recruitment website. All procedures were approved by the Mass General Brigham Institutional Review Board and took place at Mass Eye and Ear between September 2022 and February 2025. After providing informed consent, 127 study participants were screened for normal cognitive function (telephone Montreal Cognitive Assessment ≥18), English fluency (self-reported), age (18-60 years), mental health status (Beck’s Depression Inventory total score < 31), middle ear status (unremarkable otoscopy), and hearing status. Hearing status was assessed through pure tone audiometry by a licensed audiologist. Inclusion required normal audibility (≤ 25 dB HL) across the low- and mid-frequencies (0.5 up to 2 kHz) corresponding to our test stimulus and no more than mild to moderate hearing loss across the higher frequency range (3 – 8 kHz). In total, 107 study participants passed the screening criteria and participated in the study.

For this analysis, participants with constant tinnitus and notable sound sensitivity, as determined by a licensed audiologist, were excluded (n=42). Additionally, 6 participants failed to learn the task and were excluded from analyses, leaving a final sample of 59 participants that provided data for either the target-alone vs melodic distractor experiment (n = 38, 7 male) or the melodic vs. melody-matched noise distractor experiment (n = 21, 8 male). Participants completed both remote, tablet-based testing and an in-person session with EEG recording.

### 2.2 Stimuli

Target, melodic distractor, and melody-matched noise distractor stimuli were generated with pre-compensation for transducer response properties, allowing for equal-level output across frequencies. Examples of the stimuli used are available as supplementary material.

#### 2.2.1 Random or repeating target stimulus

For the laboratory-based EEG task, target stimuli consisted of concatenated sinusoidally amplitude modulated (SAM) tones (516.8 Hz carrier frequency). Five SAM rates were used to produce all random or patterned sequences: 27, 41, 54, 68, and 82 Hz. The duration of the individual SAM tones was set to 143 ms (corresponding to 6.8 Hz) to ensure that each SAM token had completed an integer number of AM and carrier frequency periods at the point of concatenation. A sequence of each SAM tokens could then either be repeated to produce a pattern segment, or a new sequence could be pseudo-randomly selected for each cycle to form random segments (restricting selection to avoid repeating the last SAM tone of the previous cycle as the first SAM tone of the next cycle).

#### 2.2.2 Melodic distractor

Melodic distractor stimuli were generated by the Magenta RL Tuner (Jaques *et al*., 2017). The output of the Magenta RL Tuner is a series of note and rest durations and corresponding note heights. This output was transposed into the frequency range below the target stimulus, maintaining a 1/3 octave protected band below the sideband of the 82 Hz SAM tone to minimize energetic masking of the target stimulus (Kidd *et al*., 2002). This melody was then duplicated 3 octaves higher, producing the identical melody in a frequency range above the target stimulus, again maintaining a 1/3 octave band from the 82 Hz SAM sideband. As per the target stimuli, notes in the melody were also amplitude modulated; note durations and SAM were adjusted to the hardware specific to remote or lab-based testing in order to produce an integer number of SAM periods per note.

#### 2.2.3 Melody-matched noise distractor

A noise stimulus was synthesized from the frequency components present in the notes of a melody to be matched; individual components lasted the entire duration of the stimulus and began with random phase. The amplitude of the frequency components was scaled to the proportion of the melody to be matched that contained that component. The imposed envelope of the melody to be matched (rest periods with zero amplitude and SAM during note periods) was then applied to the noise stimulus.

### 2.3 Psychophysical assessments of distraction

#### 2.3.1 Temporal pattern classification: speed vs accuracy task

Participants completed a speeded discrimination task in which they were asked to respond whether the target stimulus on each trial formed a pattern or random arrangement. Target stimuli lasted 4 cycles, approximately 2.9 s. A game mechanic was employed, to assess speed vs accuracy tradeoffs. A score counter hidden from view counted down from 1000 from the start of audio playback. When participants responded, they either gained (if correct) or lost (if incorrect) the points left on the counter and the score from that trial was displayed on screen. If participants failed to respond within 1 second after stimulus presentation concluded, participants lost 350 points. Blocks of trials with the target alone and the target paired with melodic distractor stimuli at different levels (18-0 dB SNR at 6 dB steps) were randomly interleaved. For the melodic distractor vs noise distractor experiment, the blocks consisted of the target alone, the target paired with melodic distractor stimuli at 12 dB SNR, and the target paired with the matched noise distractor stimuli at 12 dB SNR. Testing was self-directed and performed remotely via with a tablet computer (Microsoft Surface Pro 2, Pro 7, Redmond, WA) and calibrated closed-back circumaural headphones (Sennheiser HD280, Wedemark, Germany).

Participants underwent several stages of task familiarization before psychophysical data collection and were required to complete 50 practice trials at each phase or score ≥ 70% correct on any 10-trial block, to advance to the next familiarization stage. In practice stage 1., participants were familiarized with categorizing pattern vs random sequences of the SAM target sound. In stage 2, participants performed the temporal pattern classification task in the presence of the distractor at 18 dB SNR and then 12 dB SNR. In practice stage 3, participants were familiarized with the speeded reaction time test format described above.

#### 2.3.2 Temporal pattern classification: laboratory-based accuracy task

After completing at-home testing, participants came for a lab-based EEG recording session. The lab-based task used the same stimuli as the speed vs reaction task but was designed to minimize motion artifacts during stimulus presentation. The laboratory-based task target stimulus was always 12 cycles (∼9 s) in duration, with the first four cycles in a random arrangement and the subsequent 8 cycles were either in a random or patterned arrangement. Polarity of the stimulus alternated between cycles. Participant responses were recorded via a touchscreen tablet (Microsoft Surface Pro 4, Redmond, WA), where the virtual response buttons were provided once stimulus presentation for a given trial was complete. Auditory stimuli were delivered bilaterally through insert ear headphones (EarTone 3A, Oaktree Products, Chesterfield, MO).

Participants in the target-alone vs melodic distractor experiment completed one block with the target stimuli alone, and one block with the target and the distractor melodies at 12 dB SNR. For the melodic vs. melody-matched noise distractor experiment, the block with target alone was replaced by a block with the target and matched noise distractor as 12 dB SNR.

### 2.4 EEG Processing and Analysis

We recorded 64-channel EEG (BioSemi ActiveTwo system, Wilmington, NC), along with electrodes on the left and right mastoid, lateral to the lateral canthus of each eye, and underneath each eye. The raw EEG data were imported into the EEGLAB data structure for analysis in MATLAB (MathWorks, Natick, MA). The data were filtered between 1 and 3000 Hz using zero-phase Butterworth filters and re-referenced to the average of the left and right mastoids. Particularly noisy channels were identified using channel statistics as implemented in the FASTER pipeline (Nolan, Whelan and Reilly, 2010) and excluded from the rest of the analysis.

Epochs were individualized for each level of synchronization. For FFR analysis, cycle-length epochs time-locked to the start of each cycle were extracted. For EFR analysis, token-length epochs were taken time-locked to the start of the corresponding SAM tokens. Epochs for envelope change following response (ECFR) analysis were two-token lengths centered on the start of every other token (to avoid overlapping epochs). Regardless of window length, the phase-locking value (PLV) was calculated for each channel across epochs (subtracting the negative polarity from the positive polarity PLV for FFR analysis), and the root mean square was taken across all channels included in the analysis (Zhu *et al*., 2013). The average value of the PLV in frequency bins unrelated to the stimuli was taken as the noise floor.

The noise floor of the PLV is numerically bounded by the number of trials (Zhu *et al*., 2013). The comparisons of ECFR by magnitude of change in AM rate (Figure 1G) and for various following rates by trial response accuracy (Figure 6C-D) consisted of comparisons between conditions with different numbers of trials. To keep noise floors consistent, conditions with greater numbers of trials were randomly subsampled across 600 bootstrap iterations to the number of trials in the condition with the fewest trials.

**Figure 1.**
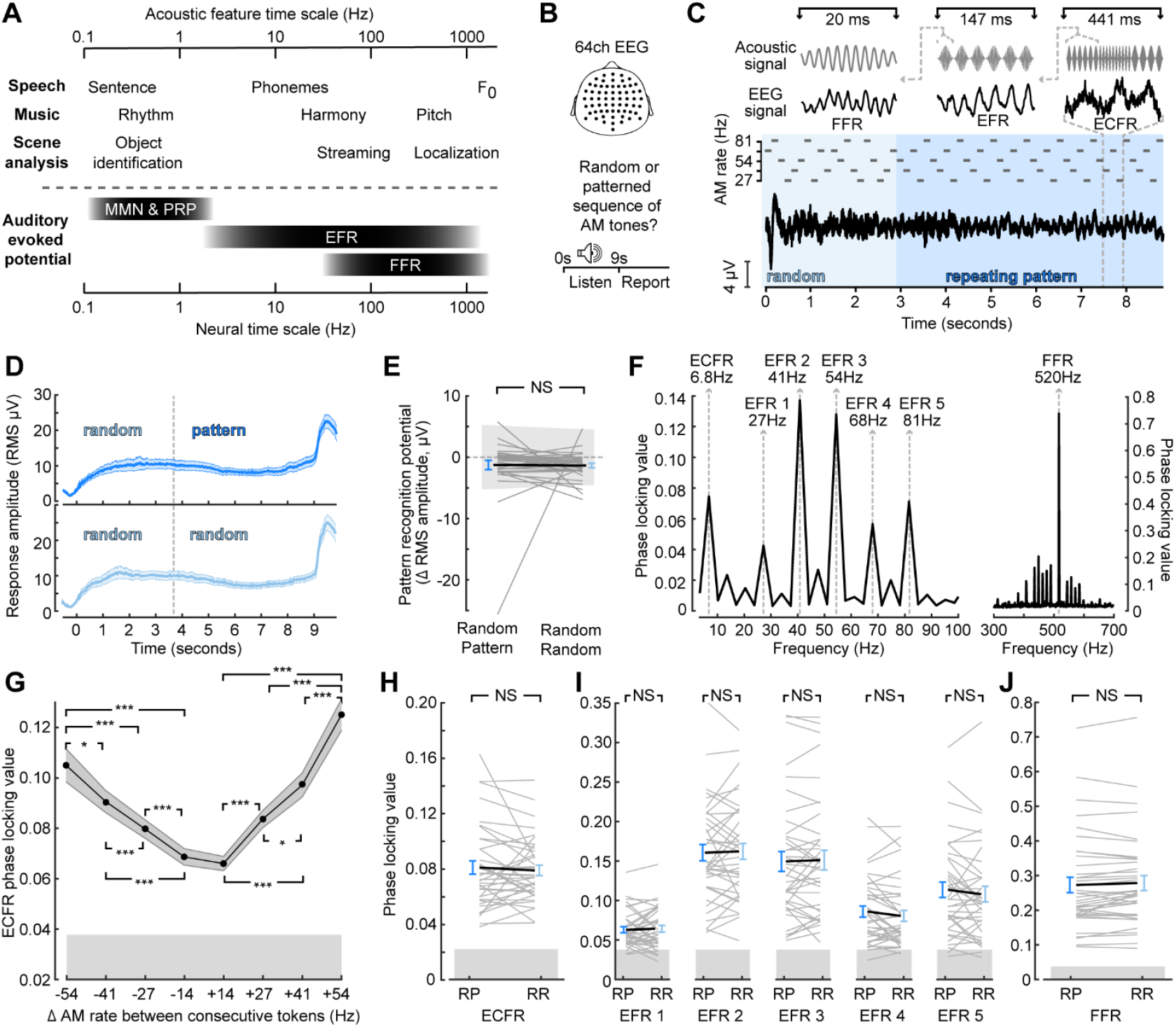
Synchronization across timescales is unaffected by target context. **(A)** Comparison of temporal features of ecologically relevant sounds (top) and EEG signals (bottom) at concomitant timescales. MMN—mismatched negativity; PRP—pattern recognition potential **(B)** 64-channel EEG was collected as participants listened to a stimulus and made a perceptual report whether the stimulus was random throughout or switched to a repeating pattern. **(C)** Illustration of the stimulus, which consisted of a sequence of SAM tones with a carrier frequency of 517 Hz, and corresponding event- related potentials. Insets at top illustrate stimulus features (left to right: carrier frequency, amplitude envelope, and changes between AM rates) and corresponding averaged EEG waveforms (left to right: FFR, EFR, ECFR). **(D)** Sustained responses to random-pattern (top, dark blue) and random-random (bottom, light blue) stimuli taking the RMS across channels **(E)** Quantification of pattern recognition potential by subtracting the two cycles before the effective transition point of random-pattern stimuli from the two cycles following the transition point for individual subjects (thin, gray) and group mean (thick, black) to random-pattern (RP) and random-random (RR) stimuli **(F)** Phase locking value spectrum of an example subject with spectral peaks corresponding to the identified feature of the target sound. **(G)** Envelope change following response (ECFR) amplitude scales with the difference in AM rate between two adjacent tokens. **(H-J)** Envelope change following responses (H), envelope following responses (I), and frequency following responses (J) are insensitive to random-pattern (RP) versus random-random (RR) context. Error bars represent the standard error of the mean. NS - not significant (*p* > 0.05); * - *p* < 0.05; *** - *p* < 0.001

We measured the sustained pattern recognition response following the procedure outlined by Southwell and colleagues (2017). Using the FieldTrip Toolbox, we filtered the data between 0.1 and 110 Hz (fifth order Butterworth filters), downsampled the data to 256 Hz, re-referenced to the average of all EEG channels, and subtracted the 1 s prestimulus baseline from the trial epoch. We then rejected trials where the average power exceeded the across-trial mean average power by two standard deviations, low-pass filtered the remaining data at 30 Hz (fifth order Butterworth filter), and applied denoising source separation for trial related activity, keeping the first five components. The sustained pattern response was the root mean square across channels of the resulting average across trials. We quantified whether the sustained response was sensitive to the pattern by subtracting the mean of the sustained response from the two target cycles before the transition point from the mean of the sustained response from the third and fourth target cycles after transition (the second and third repeats of the pattern). The same time points were used to evaluate the sustained response in the random condition. We also estimated a noise floor for this difference by bootstrapping the difference observed with a random timepoint selected as the nominal transition point.

### 2.5 Auditory Periphery Modelling

To model responses in the auditory periphery, we utilized the model published by Verhulst and colleagues (2018), version 1.2. Structures represented in the model include the middle ear, basilar membrane, inner hair cells, auditory nerve fibers (at various thresholds and spontaneous rates), the cochlear nucleus, and inferior colliculus. Model parameters were set to their defaults, and inputs to the model were an example target stimulus, the target stimulus with a melodic distractor at 12 dB signal-to-noise ratio (SNR), and the target stimulus with the corresponding matched noise distractor at 12 dB SNR.

### 2.6 Statistical Analysis

Statistical comparisons were made via t-tests, paired t-tests, and repeated measures analysis of variance (rmANOVA). Tests were done in MATLAB using the functions ttest (one-sample and paired t-tests), ttest2 (two-sample t-tests), and ranova (rmANOVA), with the function multcomp used for pairwise comparisons following rmANOVA. We control the family-wise error rate using the Bonferroni-Holm method for tests making comparisons between the same conditions. To meet the assumptions of parametric statistical testing, accuracy scores in the behavioral temporal pattern classification task were converted to rationalized arcsine units (Studebaker, 1985). For ease of comprehension, the untransformed percent correct values are reported in the text.

## 3 Results

### 3.1 Target Context and Synchronization

Target stimuli were designed to produce synchronization to acoustic features at nested timescales, similar to ethologically relevant stimuli while producing distinct neural signatures that can be measured in the frequency range with EEG (Figure 1A-B). Target stimuli acoustic features included a constant carrier frequency; SAM at one of five different rates; and changes between SAM rates at regular time intervals (Figure 2C). Represented in the EEG, these features produce an FFR, EFRs, and a waveform that follows the changes between SAM rates. SAM tones were also arranged into either patterned or random segments expected to produce a sustained pattern recognition potential that would be greater when the target context was patterned than when it was random (Barascud *et al*., 2016; Southwell *et al*., 2017; Herrmann and Johnsrude, 2018; Herrmann, Buckland and Johnsrude, 2019). While we were able to observe sustained potentials for both types of stimuli (Figure 1D), the sustained potential after transition to a patterned context did not differ from the sustained potential before the transition, and no difference was observed between random-pattern and random-random stimuli (Figure 1E; paired sample t-test *p* = 0.93).

**Figure 2.**
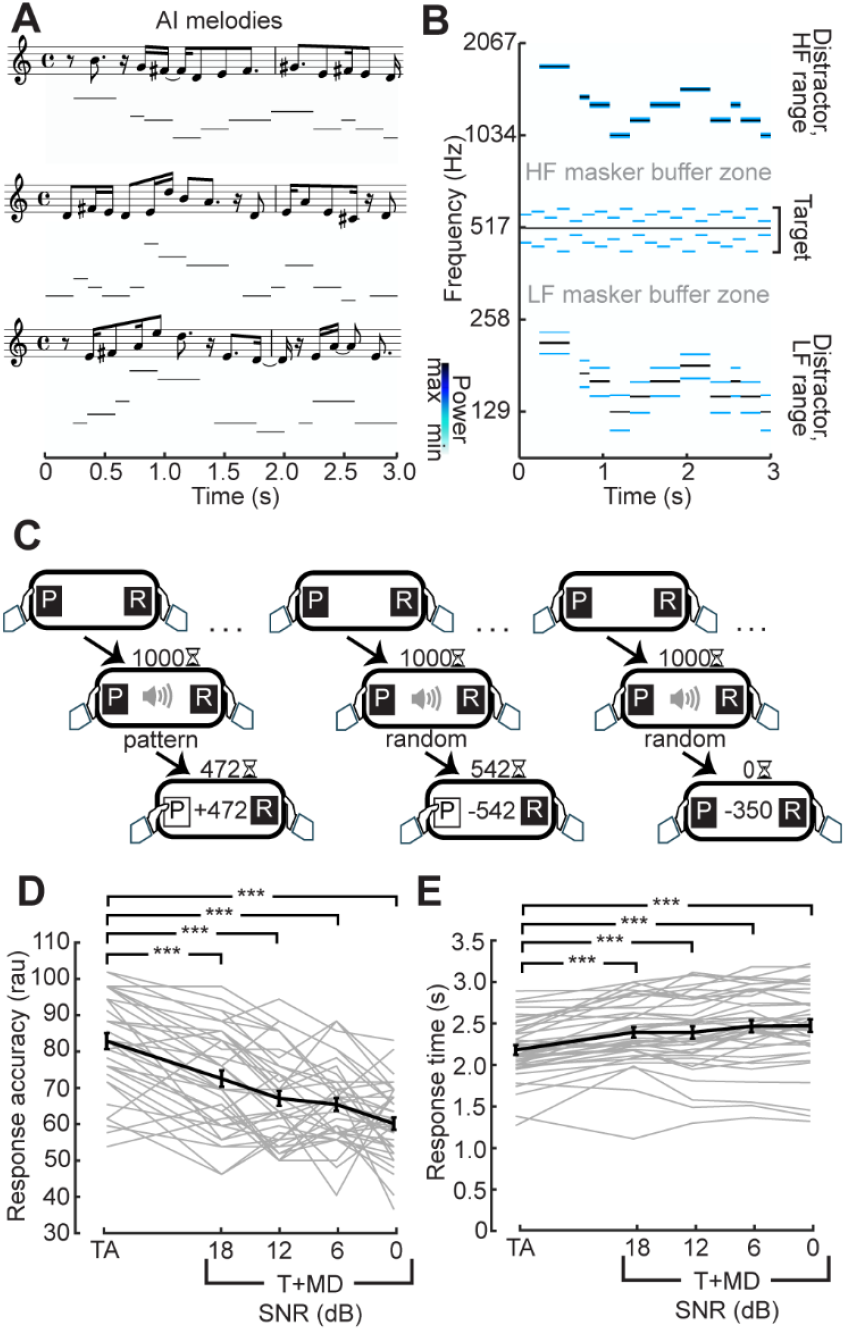
Distraction by AI-generated melodies. **(A)** Three exemplar melody sequences generated from the Magenta RL Tuner in musical notation and corresponding spectrogram representation. **(B)** Spectrogram illustrating an exemplar melody transposed above and below the target stimulus, conserving a 1/3 octave buffer zone surrounding the target to minimize energetic masking of the target. Notes in the melody are amplitude modulated at 19.5 Hz producing the sidebands seen in the spectrogram. **(C)** Temporal classification task: Participants were instructed to respond once they were confident whether the target was random or repeating. Correct responses (left) were rewarded with the points remaining on the timer whereas incorrect responses had the points subtracted from their total (middle). Trials where participants did not respond within 1.5 s after stimulus presentation was completed (right) lost participants 350 points. **(D, E)** Monotonic reduction in accuracy (D) in rationalized arcsine units and increased reaction time (E) with progressively less favorable signal-to- noise ratio (SNR). Individual subjects are shown as thin gray lines, with the group mean shown as a thick black line. Error bars reflect the standard error of the mean. *** - *p* < 0.001

To quantify the synchronization in the EEG to the acoustic features, we utilized frequency-domain analyses of the PLV. Synchronization to target stimulus features was observed for the carrier frequency (FFR) as well as each SAM rate (EFRs; Figure 1F). Additionally, we observed a peak at the rate of changing between different envelopes at 6.8Hz, which we termed the envelope change following response (ECFR). The ECFR was sensitive to the degree of AM rate difference between consecutive tokens (Figure 1G). Repeated measures ANOVA showed a significant main effect of AM difference (*F*(7, 259) = 54.94; *p* < 0.001, Greenhouse-Geisser corrected) and pairwise Tukey HSD comparisons showed statistically significant differences for all pairs between different magnitudes other than -54 to +41 (*p* = 0.64) and -41 and +27 (*p* = 0.33).

To examine the effect of patterned or random context on synchronization to target features, we compared PLVs for each acoustic feature between the random-pattern and random-random stimuli (Figure 1 H-J). Only epochs from the last 6 stimulus cycles, corresponding to after two cycles of the pattern potion of the random-pattern stimuli, were included in these analyses. Paired t-tests with Bonferroni-Holm correction showed no significant differences in ECFR (*p*-adj. = 1), EFR at 27 Hz (*p*-adj. = 1), 41 Hz (*p*-adj. = 0.75), 54 Hz (*p*-adj. = 1), 68 Hz (*p*-adj. = 0.56), and 82 Hz (*p*-adj. = 1), nor the FFR (*p*-adj. = 0.85) based on embedded context.

### 3.2 Distracting Effects on Behavior

While stimulus context (random vs patterned) had no measurable effect on EEG following responses across these various timescales, we reasoned that increased attentional demands in the form of information masking or attentional selection could exert a stronger effect (Bharadwaj, Lee and Shinn- Cunningham, 2014; Schüller *et al*., 2023). To address this possibility, we paired the auditory target with distracting melodies, as perception of both melodies and the target stimuli rely on pitch perception cues (Figure 2A,B). In the case of the target stimuli, the pitch cues come from the changing SAM rates, whereas the pitch cues in the melodic distractors come from changes in carrier frequencies. Participants completed a psychophysical task that challenged them to classify a temporal pattern as random or patterned as soon as they felt confident in their response (**Figure 2C**).

Participants were fairly accurate in their judgement (mean: 82% correct), despite the novelty and complexity of the attended sound feature and with only a few cycles of repetition in which to make the judgment (**Figure 2D**). In fact, the mean response time (2.2 s) was less than the duration of 3 cycles of the stimulus: on average, participants were both making the judgment and effectuating the motor response after fewer than 2 repeats of the pattern cycle.

The melodic distractor successfully impaired performance on the discrimination task. Repeated measures ANOVA showed a statistically significant difference in performance across the conditions both in terms of accuracy (*F*(4,148) = 44.82; *p* < 0.001) and response time in seconds (*F*(4, 148) = 31.73; p < 0.001, Greenhouse-Geisser corrected). Pairwise Tukey HSD comparisons showed responses to the target alone were more accurate (all *p* < 0.001) and faster (all *p* < 0.001) than with the melody at each SNR.

Similar effects on behavioral accuracy were observed during the EEG session (Figure 3A). The longer duration of the stimuli used in the EEG session and lack of time pressure on response led to overall higher accuracy (mean: 92% and 80% for target alone and with melodic distractors, respectively). The melodic distractors, however, still significantly impaired response accuracy when compared to the target alone (paired t-test; p < 0.001).

**Figure 3.**
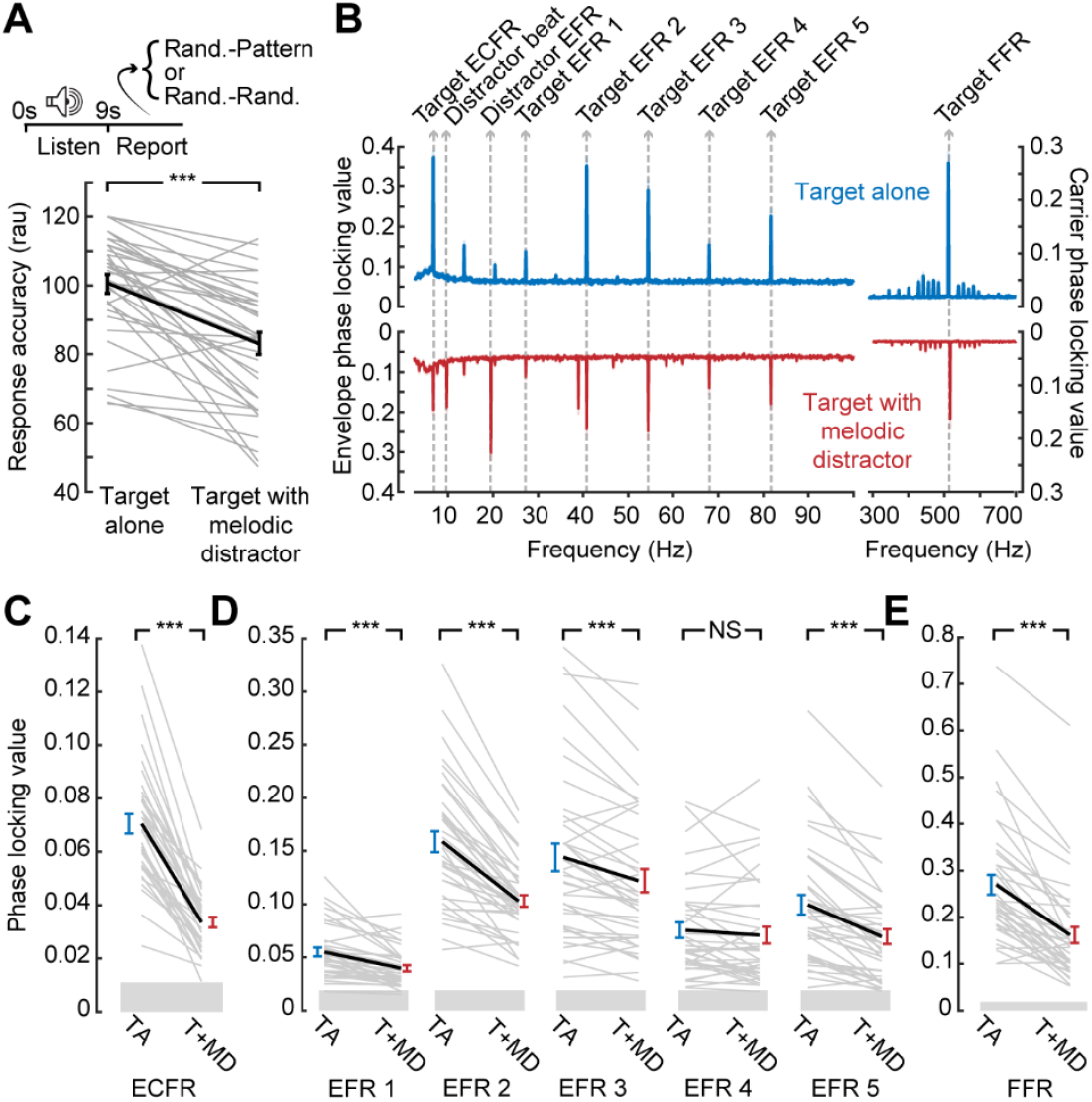
Disrupted synchronization to target sound features in the presence of a distractor. **(A)** Temporal pattern classification during the EEG session is impaired by the melodic distractor. **(B)** Grand average phase locking value spectra from the block with the target presented alone (blue) or with the melodic distractor (red). Spectral peaks corresponding to target and distractor features identified. **(C-E)** Reduced ECFR (*C*), EFR (*D*), and FFR (*E*) to the target stimuli when presented with the melodic distractor (T+MD; red) compared to the target alone (TA; blue), except for the EFR at 68 Hz. Data from individual subjects are shown as thin gray lines, with the group mean shown as a thick black line. Error bars reflect the standard error of the mean. NS – not significant (*p* > 0.05); *** - *p* < 0.001

### 3.3 Distracting Effects on Target Synchronization

EEG recordings during temporal pattern classification revealed clearly resolved FFR, EFRs, and ECFR to the target as well as synchronization peaks corresponding to the beat and envelope of the melodic distractor (Figure 3B). However, we noted a clear reduction in synchronization to all three features of the target stimulus when it was presented alongside the melodic distractor (Figure 3C-E). The ECFR PLV was decreased from 0.070 to 0.033 (*p*-adj. < 0.001); EFR PLVs decreased from 0.055 to 0.040 at 27 Hz (*p*-adj. < 0.001), from 0.16 to 0.10 at 41 Hz (*p*-adj. < 0.001), from 0.14 to 0.12 at 54 Hz (*p*-adj. < 0.001) and from 0.10 to 0.07 at 82 Hz (*p*-adj. < 0.001); and the FFR decreased from 0.27 to 0.16 (*p*-adj. < 0.001). The EFR at 68 Hz was not significantly different, with a mean target alone PLV of 0.076 and a mean target with distractor PLV of 0.071 (*p*-adj. = 0.23).

### 3.4 Controlling for Peripheral Effects of Melodic Distractors

Reduced neural encoding of the target sound in the presence of the melody could be a neurophysiological signature of distraction. Alternatively, it could reflect destructive interference between the neural signals generated by the target and distractor sounds. To better understand the underlying source of the reduced target synchronization, we utilized a biophysical model of the auditory periphery and early central pathway designed to produce modeled outputs up to the level of the auditory brainstem, including EFRs (**Figure 4A-B**; Verhulst, Altoè and Vasilkov, 2018).

**Figure 4.**
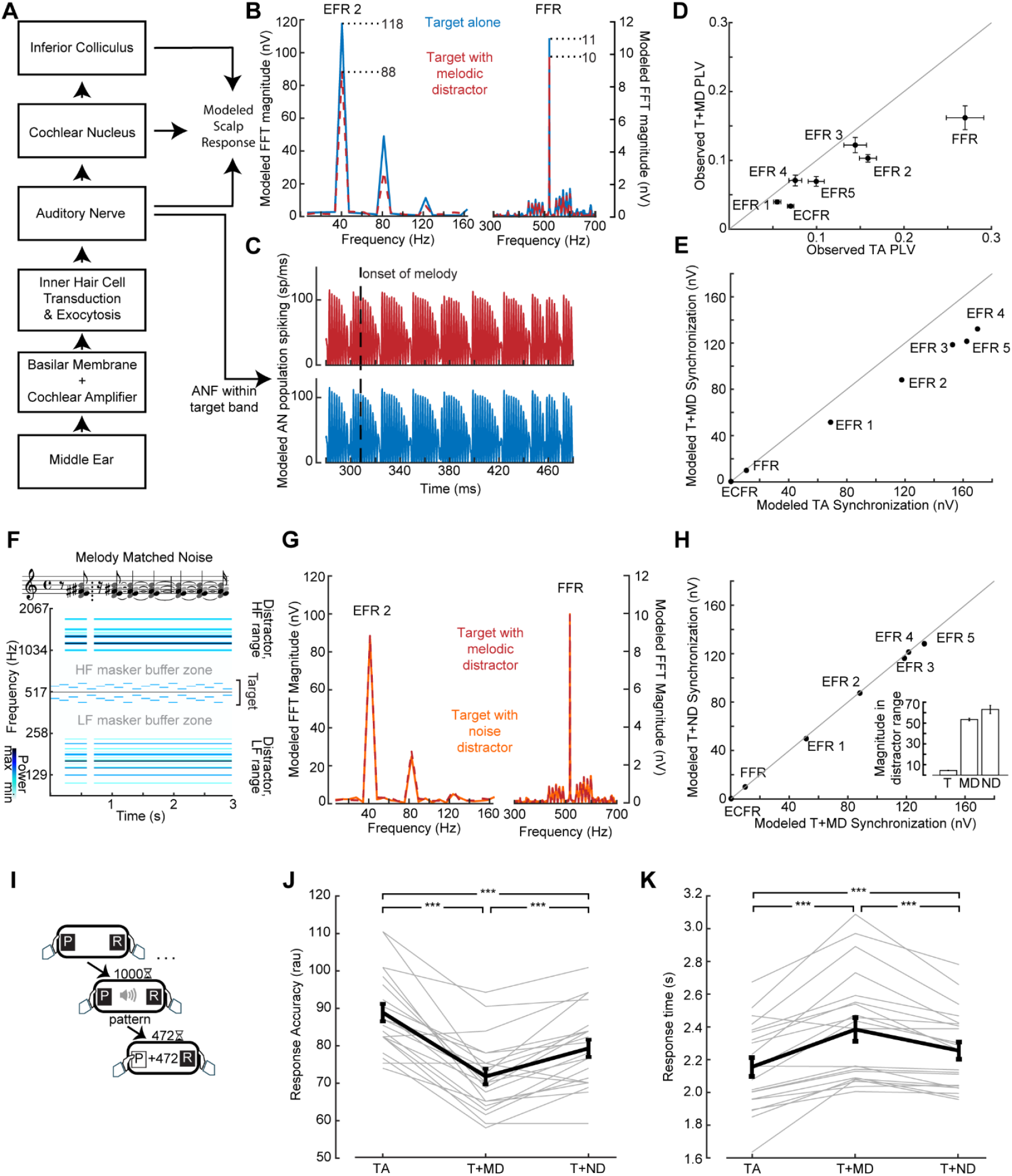
Validating distractor stimuli via model-based simulations of neural synchronization. **(A)** Simplified representation of the biophysical model from Verhulst, Altoè and Vasilkov (2018)Verhulst, Altoè and Vasilkov (2018)Verhulst, Altoè and Vasilkov (2018) used to evaluate peripheral responses to the stimuli used in this study. **(B, C)** Simulated spectra of potentials at the scalp (*B*) and responses in the auditory nerve (*C*) show weaker envelope and frequency following responses when the target is submitted to the model with the melodic distractor (red) compared to the target alone (blue) despite conserved spiking in the modeled auditory nerve. **(D, E)** FFR and EFR differences of target-alone vs target + distractor conditions measured in vivo (*D*) and simulated by the biophysical model (*E*). **(F)** Matched noise stimuli are made by including the frequencies present in the melodies (represented as notes in musical notation) across the entire duration of the stimulus, conserving “rest” periods of silence. The corresponding spectrogram of the matched noise stimulus with a target stimulus is also shown. **(G)** Simulated scalp potential spectra show similar envelope and frequency following responses when the target is submitted to the model with the melodic distractor (red) and the matched noise distractor (orange). **(H)** The effect shown in **(G)** is consistent across envelopes (points lie on the line of unity); inset: total magnitude of the spectrogram outside of the target buffer zone for the target (T), melodic distractor (MD) and matched noise distractor (ND) shows the energy was conserved between melodic and matched noise distractors. **(I)** The temporal pattern classification task, as per Figure 2. **(J-K)** Behavioral performance measured by response accuracy (*J*) and response time (*K*) is at an intermediate level when the target is paired with the noise distractor (T+ND) compared to the target alone (TA) and target with the melodic distractor (T+MD). Data from individual subjects are shown as thin gray lines, with the group mean shown as a thick black line. Error bars reflect the standard error of the mean. *** - *p* < 0.001

Population responses from modeled auditory nerve fibers with characteristic frequencies within the target stimulus bandwidth were mostly unaffected by the inclusion of the melodic distractor in the input to the model (**Figure 4C**). At the level of the simulated EFR, however, synchronization to the target carrier frequency and target envelopes were reduced (Figure 4B, E) similar to our in vivo results (replotted in **Figure 4D**). The ECFR was not observed in the modeled response.

To better account for the off-frequency effects and interference that contribute to reduced target synchronization in the presence of the melodic distractor, we created a set of matched noise stimuli designed to induce comparable neural interactions with the target as the melodic distractor but less perceptual interference. The matched noise stimuli are comprised of the same frequency components as individual melodic distractor stimuli, but with the energy at each frequency spread out across the duration of the stimulus (**Figure 4F**). The same SAM applied to the melodic distractor is also applied to the matched noise distractor. When the matched noise stimulus is included in the input to the model, the FFR and EFRs to the target stimulus match the modeled responses with the melodic distractor (**Figure 4G, H**). Comparing the target + melodic distractor to the target + noise distractor, then, would control for low-level interference effects that are unrelated to distraction.

As a next step, we set out to test the hypothesis that the matched noise distractor produced an intermediate level of perceptual interference such that temporal classification would be intermediate to target alone and the melodic distractor. We addressed this hypothesis in a second cohort of study participants with the speeded reaction time behavioral assay (**Figure 4I**). We found that mean accuracy was 87% with the target alone, in line with the previous cohort. When the melodic distractor was included, mean accuracy dropped to 72% but improved with the matched noise distractor back to 79%. Repeated measures ANOVA showed a significant main effect of condition (*F*(2, 40)=42.13; *p* < 0.001), with pairwise comparisons showing significant differences between all three conditions (all *p* < 0.001; **Figure 4J**). Mean response times were 2.16 s to the target alone, 2.38 s to the target with the melodic distractor, and 2.26 s to the target with the noise distractor. Repeated measures ANOVA showed a significant main effect of condition (*F*(2, 40)=31.06; p < 0.001, Greenhouse-Giesser corrected), with pairwise comparisons showing significant differences between all three conditions (all *p* < 0.001; **Figure 4K**).

### 3.5 Differential Distraction Effect on Target Synchronization

Having demonstrated that target + melody versus target + matched noise distractor provides the means for a more direct comparison of variable distraction loads, we performed another in-lab assessment of combined neural and behavioral recordings. We confirmed that behavioral accuracy during the EEG recordings recapitulated the results from the speeded reaction time temporal classification task (93% vs. 80%; paired t-test *p* < 0.001; **Figure 5A**). When the matched noise is paired with the target stimulus, as with the melody, we observe synchronization peaks at the relevant target frequencies (**Figure 5B**). We also observe peaks corresponding to the envelope of the distractor, but the peak at the “beat” of the melody is not reproduced with the matched noise as it does not contain the note changes present in the melody.

**Figure 5.**
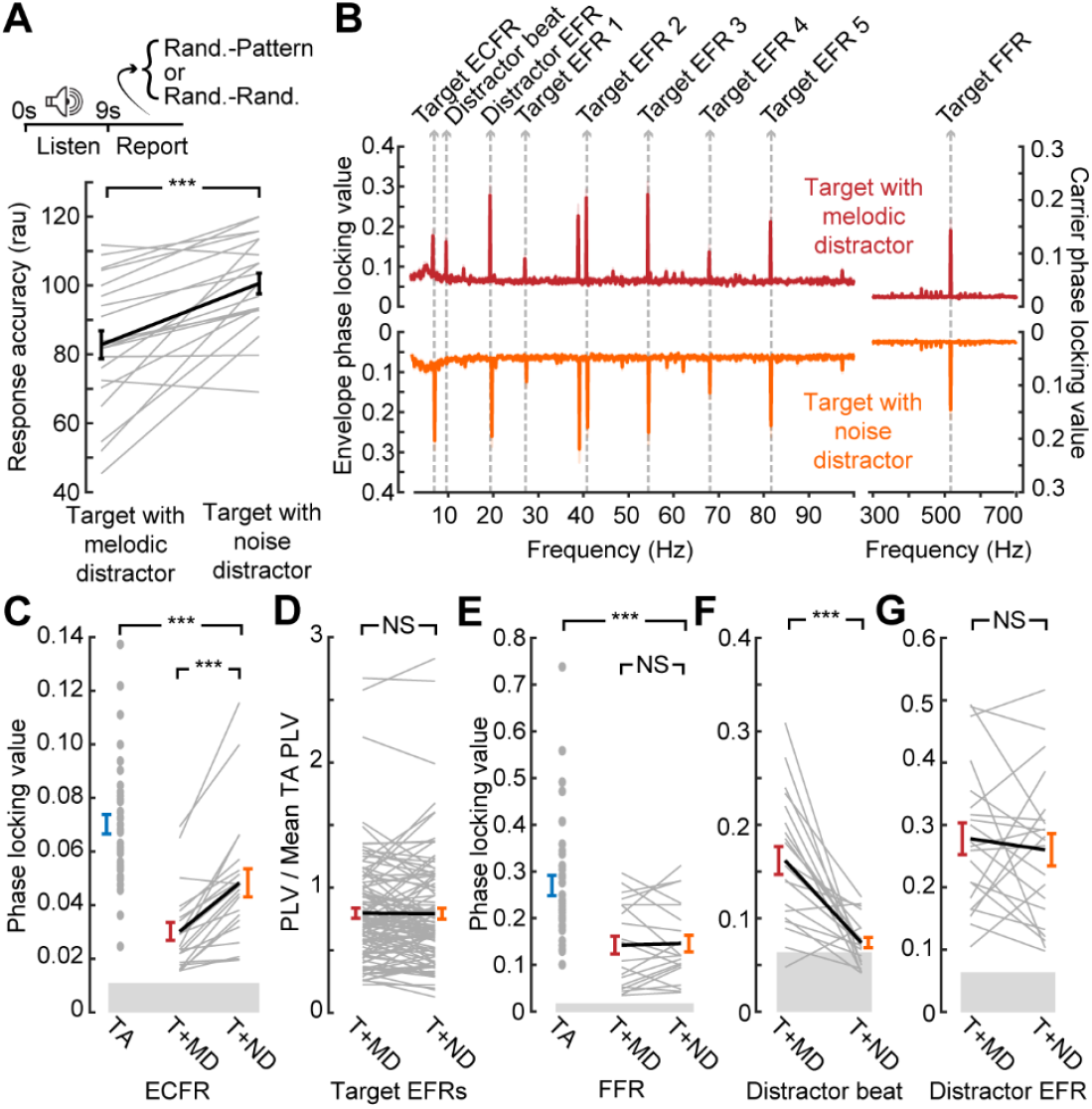
Only the slowest timescale of neural target synchronization, the ECFR, is modulated by distraction level. **(A)** Data collection paradigm, as per Figures 1 and 3. **(B)** Phase locking value spectra of responses to the target with melodic distractor (red) and the target with matched noise distractor (orange). Peaks related to target and distractor stimuli features are labeled. **(C)** ECFRs to the target with matched noise (T+ND; orange) distractor are greater than those observed for the target with melodic distractor (T+MD; red), but less than those observed for the target alone in the prior cohort (TA; blue). **(D, E)** EFRs, normalized to the mean target alone PLV and collapsed across AM rate (*D*), and FFRs (*E*) do not show significant differences between the melodic distractor and noise distractor conditions. The FFR is reduced relative to the target alone in the prior cohort (TA; blue). **(F)** The matched noise distractor does not reliably produce synchronization at the melodic beat rate. **(G)**Synchronization to distractor envelope does not differ by distractor type. Data from individual subjects are shown as thin gray lines, with the group mean shown as a thick black line. Error bars reflect the standard error of the mean. NS – not significant (*p* > 0.05); *** - *p* < 0.001

We found that neural synchronization to the target ECFR was significantly greater in the less distraction matched-noise condition than the melody block (mean 0.048 vs 0.030; *p*-adj. < 0.001; **Figure 5C**). Additionally, only two individual subjects went against the group trend. A two-sample t-test also showed that the ECFR in the matched noise block in the second cohort was significantly lower than that observed with the target alone in the prior cohort (*p*-adj. < 0.001).

While the EFR strength varied across different AM rates, the effects of different conditions appeared to be consistent. We therefore consolidated EFRs across AM rate by normalizing the values to the mean target alone values for each AM rate (Figure 5D). The resulting normalized values showed no difference between EFRs in the melody compared to the matched noise (*p*-adj. = 0.86). The FFR (Figure 5E) also showed no difference between the melody and the matched noise blocks (*p*-adj. = 1), but a two-sample t-test comparing the matched noise FFR to the target alone FFR in the prior cohort did show a significant effect (*p*-adj. < 0.001).

We also examined synchronization to the distractor stimuli at the melody beat rate (Figure 5F) and the distractor SAM rate (Figure 5G). As shown in the spectra, synchronization to the melody beat rate was only clearly present when the melody was presented and approached the noise floor when the matched noise was presented, leading to a significant difference between conditions (*p*-adj. < 0.001). No differences were observed between melodic and matched distractor EFR (*p*-adj. = 1).

Therefore, of all the target stimulus features that could be reliably measured with both distractors, only the ECFR showed a differential effect of distraction between the more distracting melody condition and the less distracting matched noise condition.

### 3.6 Synchronization Changes Based on Perceptual Report

To strengthen the inference that the differential effect between the melodic and noise distractors on ECFR is due to distraction by the melodies, we examined synchronization on correct versus incorrect response trials (Figure 6A). We pooled participants from both cohorts, and examined the data from the melody block, which was common to both participant cohorts (Figure 6B). Participants with fewer than 15% error trials with the melodic distractor were excluded from this analysis (n = 24). For the remaining participants, following responses were separately calculated for correct and incorrect trials (**Figure 6C, D**). Because the number of trials, and thus the noise floor of the phase locking value (Zhu *et al*., 2013), in each trial category varies between participants, we quantified the difference in following rates in terms of an asymmetry index (correct – incorrect)/(correct + incorrect)), and only included the asymmetry index for each participant and following rate when at least one category was above the noise floor for that following rate. One-sample t-tests showed that asymmetry indices for the FFR, ECFR, and melody beat response were significantly different than zero (*p* = 0.03-, 0.041, and 0.0018, respectively). We note, however, the small effect size for the FFR. While the result is statistically significant, the mean asymmetry index is -0.026 for FFR, compared to 0.043 for ECFR and -0.11 for the melody beat response. Both target EFRs and the distractor EFR did not differ by response accuracy (*p* = 0.45, and 0.38, respectively).

**Figure 6.**
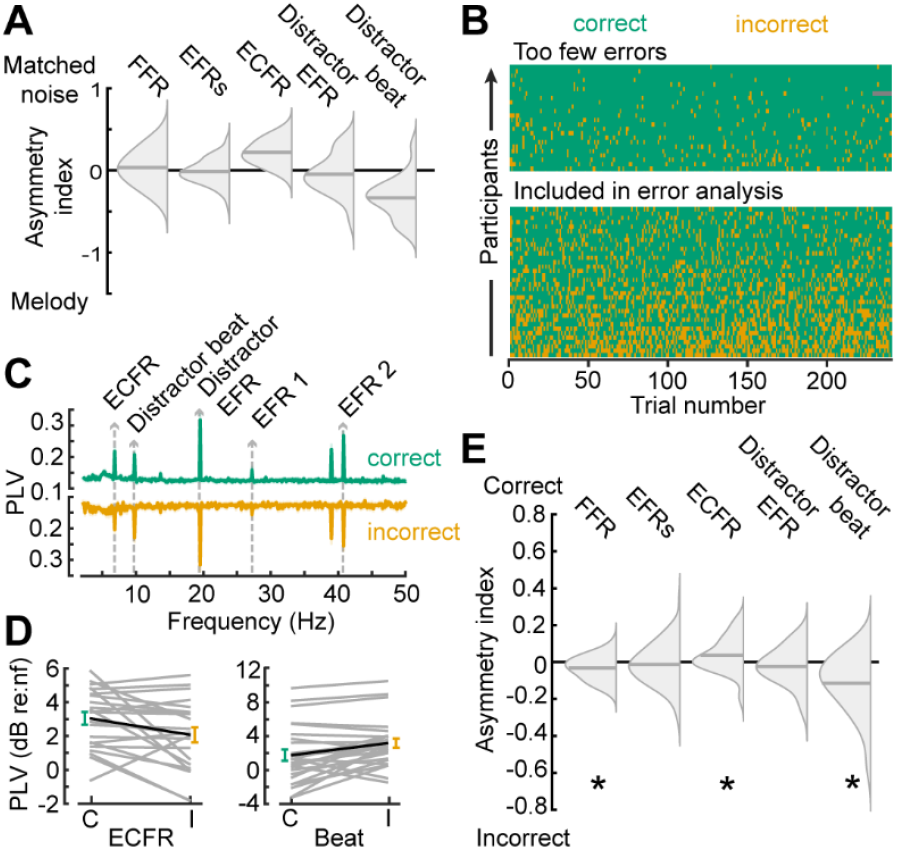
Synchronization to target changes is decreased while synchronization to distractor changes is increased during incorrect trials. **(A)** Summary of findings in Figure 5, plotted as split violin plots of the asymmetry index between the matched noise and melodic distractor conditions. **(B)** Raster of trial-by-trial response accuracy for all participants during the melodic distractor block, sorted by overall accuracy. Participants with too few error trials (top section) were excluded from analysis of the error trials. Green represents correct trials, yellow incorrect trials, and gray missing trials due to technical error. **(C)** Phase locking value spectra of correct (green, top) and incorrect (yellow, bottom) trials, focused on the lower frequency synchronization peaks. **(D)** The ECFR (*left*) and melody beat synchronization (*right*) plotted in dB relative to individual subject noise floors (nf) for correct (green, *C*) and incorrect (yellow, *I*) trials. Data from individual subjects are shown as thin gray lines, with the group mean shown as a thick black line. **(E)** Asymmetry indices for synchronization peaks between correct and incorrect trials show decreased ECFRs and increased synchronization to the melody beat during incorrect trials. FFRs are also slightly elevated during incorrect trials. * - *p* < 0.05

## 4 Discussion

Here, we developed a paradigm that allows us to measure neural synchronization across multiple timescales and with multiple stimuli presented, including synchronization to both target and competitor stimuli. This paradigm has yielded an electrophysiological measurement that is sensitive to auditory distraction which may be applied in future studies to populations of interest.

### 4.1 Synchronization Across Timescales

We have shown it is possible to measure synchronization across multiple timescales simultaneously with carefully designed stimuli. Naturalistic stimuli contain temporal dynamics across a multitude of timescales with different types of features. Past studies of synchronization in the auditory nervous system, even with complex stimuli, have typically focused on a single feature and timescale at a time (Krizman and Kraus, 2019). These studies have found important objective measures that provide insight into various auditory processes and disorders (Kraus *et al*., 2016; Vercammen *et al*., 2018; Parthasarathy *et al*., 2020; Bidelman and Momtaz, 2021) but are limited by only analyzing a single timescale. Features at different timescales may be differentially implicated in various auditory perception tasks (Swaminathan *et al*., 2016) and implicate different auditory structures (Kuwada *et al*., 2002; Coffey *et al*., 2019), so a design that gathers across multiple timescales provides advantages that single features do not, as other recent paradigms demonstrate (Wang, Bharadwaj and Shinn-Cunningham, 2019; Calcus, Undurraga and Vickers, 2022; Schüller *et al*., 2023).

We identified a novel following response, the ECFR. With our target stimulus, the ECFR is elicited by changes between different SAM envelopes which occur at a consistent rate. We also, however, observe a similar following rate to regular changes in a stimulus with the melody beat following rate (Figure 1G). This suggests that the ECFR is part of a larger class of synchronization to changes in ongoing stimuli as has been observed for inter-aural phase difference changes (Undurraga *et al*., 2016; Parthasarathy *et al*., 2020) and synchronization to syllable rate in speech stimuli (Assaneo *et al*., 2019; He, Buder and Bidelman, 2023). These synchronized responses may also be related to the acoustic change complex (Kim, 2015) traditionally elicited by infrequent changes in the stimuli. In fact, at least one other study has reported acoustic change complexes evoked by stimuli with carrier frequency changes at rates similar to the envelope changes used in our study and observed a similar synchronization peak at the rate of change (Calcus, Undurraga and Vickers, 2022).

Synchronization at all timescales was affected by the presence of a competing sound. Previous work with noise stimuli had primarily shown effects at the harmonics of the EFR (Zhu *et al*., 2013), but our analyses show significant impacts even at the fundamental of both the FFR and the EFR. This occurs despite reducing energetic masking of the target stimulus by keeping frequencies in the distractor separated from the frequencies present in the target. Using a model of the auditory periphery, we show that effects on the FFR and EFRs are partially explained by off-frequency contributors to these following rates. Importantly, the model that we used has no cortical component, despite the generators of the EFRs likely including cortical sources (Kuwada *et al*., 2002; Picton *et al*., 2003; Shinn-Cunningham, Varghese, Wang and Bharadwaj, 2017). Future studies that examine following responses when multiple sound sources are presented should take care to account for such low-level interference between stimuli, even in the absence of direct energetic masking.

### 4.2 No Effect of Pattern

Previous studies have reported a sustained potential difference related to repeating patterns in complex stimuli using MEG (Barascud *et al*., 2016) and EEG (Southwell *et al*., 2017; Herrmann and Johnsrude, 2018; Herrmann, Buckland and Johnsrude, 2019). With our target stimuli, however, we did not observe an effect of the pattern or random context. Both the pattern recognition potential and the following responses to the low-level features were unchanged by this context. Differences between the target stimuli utilized in our paradigm and previous studies include greater overall predictability in the stimulus, a potentially less-salient feature to differentiate tones, and active listening in our paradigm compared to passive listening in previous studies. The change in the sustained potential during a patterned stimulus has been associated with the predictability of the stimulus (Barascud *et al*., 2016; Southwell *et al*., 2017). Previous paradigms involved larger pools of potential combinations to select from; for example, in the work first describing this potential (Barascud *et al*., 2016), tones were selected from a library of 20 possible tones to build into sequences. While they were able to show effects when the chosen set on a given trial was limited to only 5 of the tones—in line with the five AM tones used in our target stimuli—the trial-to-trial variability of the selected tones contributed to greater uncertainty than in our paradigm. Additionally, the differences between tones in our target stimuli—changes in SAM rate—may be more subtle than changes in frequency (Barascud *et al*., 2016; Southwell *et al*., 2017) or coherence of modulation among several tones (Herrmann and Johnsrude, 2018; Herrmann, Buckland and Johnsrude, 2019) that have been used previously. Finally, previous studies have employed passive listening to the patterned stimuli, whereas here we asked participants to attend to and report on the pattern or random context of the stimuli.

### 4.3 Distraction

We characterize the effect of the melodic stimuli on target perception as distraction. More broadly, the effect falls under informational masking (Durlach, Mason, Kidd, *et al*., 2003). While informational masking is an appropriate description of how the melodic stimuli impair target stimulus perception, we avoided the intentional use of low-level acoustic features classically associated with informational masking. Timing coincidences and manipulations of the same features in both targets and maskers have previously been utilized to produce informational masking, in part through perceptual grouping (Durlach, Mason, Shinn-Cunningham, *et al*., 2003). As our approach depended on isolating neural entrainment to distinct and independent temporal features of the target and distractor sounds, we could not manipulate distraction through introducing greater temporal co- incidence between sound features. Instead, we sought to manipulate the salience of the distractor rather than manipulating low-level acoustic features to match the target stimuli.

The behavioral results, taken together, are consistent with distraction being a key feature of the impaired target perception. Participants were both less accurate and slower to respond when the target was accompanied by the melodic distractor than when the target was alone (Figure 2D,E; Figure 3A) or when the target was accompanied by the matched noise distractor (Figure 4J,K; Figure 5A). It is also notable that in the EEG session, when participants had a longer time to suppress the distractor and evaluate the target, accuracy was similar between the target alone block for the first cohort (mean 92%) and the matched noise block for the second cohort (mean 91%). The less- distracting matched noise stimuli were behaviorally completely suppressed, given enough time to do so, while the melodies continued to impair target perception.

Additionally, the trade-off between target and distractor synchronization observed in the analysis by response accuracy (Figure 6) is indicative of an attentional shift from the target to the melodic distractor (Kong, Mullangi and Ding, 2014). When participants make errors—and are thus failing to correctly perceive the target stimulus—we observe increased synchronization to the changes in the distractor and decreased synchronization to changes in the target. If the informational masking effect of the melodies was driven by failures of stream segregation, we would expect synchronization to both target and distractor to be reduced as the features of the combined stream would be ambiguous (Choi *et al*., 2014). That we instead observe a trade-off indicates that target and melodic stimuli were still segregated, but that the distractor melodies received greater attention on trials when participants made errors.

### 4.4 Low-level Synchronization Insensitive to Distraction

The FFR and EFRs were insensitive to differential levels of distraction once off-target effects of competitor stimuli were accounted for by the matched noise stimulus (Figure 5). This result is in line with previous work that has shown FFR and EFR measurements that are insensitive to attentional state (Varghese, Bharadwaj and Shinn-Cunningham, 2015). While some studies have shown an effect of attention on EFRs at frequencies around 40 Hz, these effects are typically shown with spatially separated streams (Müller *et al*., 2009; Bharadwaj, Lee and Shinn-Cunningham, 2014). Spatial segregation is a helpful auditory cue in natural listening environments, but spatial auditory attention involves different processes than those evoked under non-spatial tasks (Bonacci, Bressler and Shinn-Cunningham, 2020). The increased EFR observed in other studies may be a result of the recruitment of this spatially-selective system; our stimuli were presented diotically, thus providing no spatialized cues.

### 4.5 Neural Measure Sensitive to Distraction

We have demonstrated that the ECFR (and indirectly the melody beat following response) is sensitive to the level of auditory distraction. That synchronization to these measures occurred in or near the alpha band of activity is perhaps significant. Prior work has demonstrated a link between alpha oscillations and inhibition of distractors (Kelly *et al*., 2006; Foxe and Snyder, 2011; Händel, Haarmeier and Jensen, 2011). This work has also been extended to the auditory modality (Banerjee *et al*., 2011; Deng *et al*., 2019; Deng, Choi and Shinn-Cunningham, 2020). It is notable, however, that this link has been established primarily with regards to spatial attention, and there were no spatial cues provided in our paradigm. Furthermore, while alpha oscillations related to distractor suppression have been shown to be linked to stimulus-related synchronous activity at lower frequencies (Wöstmann *et al*., 2016), the relationship between stimulus-related synchronous activity within the alpha band and top-down distractor suppression alpha activity has not yet been explored.

Susceptibility to auditory distraction is implicated in a variety of conditions, including difficulty understanding speech in noisy environments (Shinn-Cunningham, amplification and 2008, 2008; Tierney, Rosen, Psychology, *et al*., 2020), tinnitus (McKenna *et al*., 2014), and neurodevelopmental disorders(O’Connor, 2012; Blomberg *et al*., 2019; Keehn *et al*., 2019). While the ECFR cannot serve as a direct measure of these disorders, it may be useful in evaluating this contributing factor.

Moreover, there is evidence that auditory distractor suppression may be a skill that can be trained through interactive training software (Whitton *et al*., 2017; de Larrea-Mancera *et al*., 2022).

Measurement of factors that are receptive to change may prove particularly useful in evaluating therapeutic effects. Future studies may evaluate longitudinal changes in the ECFR as participants receive training in suppressing auditory distractors. Individuals who improve in their ability to suppress distractors may see increases in ECFR when target stimuli are accompanied by highly distracting competitors.

The ECFR, but not the FFR or EFRs, is a neural measure sensitive to the level of auditory distraction. Being measured alongside the low-level, distraction-insensitive FFR and EFRs provides an internal control for generalized disordered temporal processing. Our results point to the potential of paradigms evoking the ECFR and other regular stimulus changes to elicit objective measures of auditory distraction in a variety of contexts.

## 6 Author Contributions

DS: Conceptualization, Formal analysis, Methodology, Software, Visualization, Writing—original draft. JS: Data curation, Investigation, Methodology, Resources, Writing—review & editing. AP: Methodology, Writing—review & editing. KH: Resources, Software, Writing—review & editing. DP: Conceptualization, Funding acquisition, Project administration, Supervision, Visualization, Writing—review & editing.

## 8 Acknowledgments

We thank Gerald Kidd, Hari Bharadwaj, Josh McDermott, and Barbara Shinn-Cunningham for feedback that motivated the development of the matched noise stimulus. Finally, we thank all the participants who agreed to undergo study procedures in order to provide the data described herein.

## 10 Data Availability Statement

The raw data supporting the conclusions of this article and code used to analyze the data will be made available by the authors, without undue reservation.

## References

Assaneo, M.F., Ripollés, P., Orpella, J., Lin, W.M., de Diego-Balaguer, R. and Poeppel, D. (2019) ‘Spontaneous synchronization to speech reveals neural mechanisms facilitating language learning’, Nature Neuroscience, 22(4), pp. 627–632. Available at: 10.1038/s41593-019-0353-z.

Banerjee, S., Snyder, A.C., Molholm, S. and Foxe, J.J. (2011) ‘Oscillatory alpha-band mechanisms and the deployment of spatial attention to anticipated auditory and visual targetlocations: Supramodal or sensory-specific control mechanisms?’, Journal of Neuroscience, 31(27), pp. 9923–9932. Available at: 10.1523/JNEUROSCI.4660-10.2011.

Barascud, N., Pearce, M.T., Griffiths, T.D., Friston, K.J. and Chait, M. (2016) ‘Brain responses in humans reveal ideal observer-like sensitivity to complex acoustic patterns’, Proceedings of the National Academy of Sciences of the United States of America, 113(5), pp. E616–E625. Available at: 10.1073/PNAS.1508523113.

Bharadwaj, H.M., Lee, A.K.C. and Shinn-Cunningham, B.G. (2014) ‘Measuring auditory selective attention using frequency tagging’, Frontiers in Integrative Neuroscience, 8(FEB). Available at: 10.3389/FNINT.2014.00006/FULL.

Bharadwaj, H.M., Masud, S., Mehraei, G., Verhulst, S. and Shinn-Cunningham, B.G. (2015) ‘Individual Differences Reveal Correlates of Hidden Hearing Deficits’, The Journal of Neuroscience, 35(5), p. 2161. Available at: 10.1523/JNEUROSCI.3915-14.2015.

Bharadwaj, H.M. and Shinn-Cunningham, B.G. (2014) ‘Rapid acquisition of auditory subcortical steady state responses using multichannel recordings’, Clinical neurophysiology, 125(9), p. 1878. Available at: 10.1016/J.CLINPH.2014.01.011.

Bidelman, G.M. and Momtaz, S. (2021) ‘Subcortical rather than cortical sources of the frequency-following response (FFR) relate to speech-in-noise perception in normal-hearing listeners’, Neuroscience Letters, 746. Available at: 10.1016/j.neulet.2021.135664.

Blomberg, R., Danielsson, H., Rudner, M., Söderlund, G.B.W. and Rönnberg, J. (2019) ‘Speech Processing Difficulties in Attention Deficit Hyperactivity Disorder’, Frontiers in Psychology, 10, p. 458190. Available at: 10.3389/fpsyg.2019.01536.

Bonacci, L.M., Bressler, S. and Shinn-Cunningham, B.G. (2020) ‘Nonspatial features reduce the reliance on sustained spatial auditory attention’, Ear and Hearing, 41(6), pp. 1635–1647. Available at: 10.1097/AUD.0000000000000879.

Brungart, D.S., Simpson, B.D., Ericson, M.A. and Scott, K.R. (2001) ‘Informational and energetic masking effects in the perception of multiple simultaneous talkers’, The Journal of the Acoustical Society of America, 110(5), pp. 2527–2538. Available at: 10.1121/1.1408946.

Calcus, A., Undurraga, J.A. and Vickers, D. (2022) ‘Simultaneous subcortical and cortical electrophysiological recordings of spectro-temporal processing in humans’, Frontiers in Neurology, 13. Available at: 10.3389/FNEUR.2022.928158/FULL.

Cancel, V.E., McHaney, J.R., Milne, V., Palmer, C. and Parthasarathy, A. (2023) ‘A data-driven approach to identify a rapid screener for auditory processing disorder testing referrals in adults’, Scientific Reports, 13(1), p. 13636. Available at: 10.1038/s41598-023-40645-0.

Choi, I., Wang, L., Bharadwaj, H. and Shinn-Cunningham, B. (2014) ‘Individual differences in attentional modulation of cortical responses correlate with selective attention performance’, Hearing Research, 314, pp. 10–19. Available at: 10.1016/j.heares.2014.04.008.

Coffey, E.B.J., Nicol, T., White-Schwoch, T., Chandrasekaran, B., Krizman, J., Skoe, E., Zatorre, R.J. and Kraus, N. (2019) ‘Evolving perspectives on the sources of the frequency-following response’, Nature Communications, 10(1), p. 5036. Available at: 10.1038/s41467-019-13003-w.

Cooke, M., Garcia Lecumberri, M.L. and Barker, J. (2008) ‘The foreign language cocktail party problem: Energetic and informational masking effects in non-native speech perception’, The Journal of the Acoustical Society of America, 123(1), pp. 414–427. Available at: 10.1121/1.2804952.

Deng, Y., Choi, I. and Shinn-Cunningham, B. (2020) ‘Topographic specificity of alpha power during auditory spatial attention’, NeuroImage, 207(November 2019), p. 116360. Available at: 10.1016/j.neuroimage.2019.116360.

Deng, Y., Reinhart, R.M.G., Choi, I. and Shinn-Cunningham, B. (2019) ‘Causal links between parietal alpha activity and spatial auditory attention’, eLife, 8, pp. 1–23. Available at: 10.7554/eLife.51184.

Durlach, N.I., Mason, C.R., Kidd, G., Arbogast, T.L., Colburn, H.S. and Shinn-Cunningham, B.G. (2003) ‘Note on informational masking (L)’, The Journal of the Acoustical Society of America, 113(6), pp. 2984–2987. Available at: 10.1121/1.1570435.

Durlach, N.I., Mason, C.R., Shinn-Cunningham, B.G., Arbogast, T.L., Colburn, H.S. and Kidd, G. (2003) ‘Informational masking: Counteracting the effects of stimulus uncertainty by decreasing target-masker similarity’, The Journal of the Acoustical Society of America, 114(1), pp. 368–379. Available at: 10.1121/1.1577562.

Foxe, J.J. and Snyder, A.C. (2011) ‘The role of alpha-band brain oscillations as a sensory suppression mechanism during selective attention’, Frontiers in Psychology, 2(JUL), pp. 1–13. Available at: 10.3389/fpsyg.2011.00154.

Galambos, R., Makeig, S. and Talmachoff, P.J. (1981) ‘A 40-Hz auditory potential recorded from the human scalp’, Proceedings of the National Academy of Sciences of the United States of America, 78(4 II), pp. 2643–2647. Available at: 10.1073/PNAS.78.4.2643.

Händel, B.F., Haarmeier, T. and Jensen, O. (2011) ‘Alpha oscillations correlate with the successful inhibition of unattended stimuli’, Journal of Cognitive Neuroscience, 23(9), pp. 2494–2502. Available at: 10.1162/jocn.2010.21557.

He, D., Buder, E.H. and Bidelman, G.M. (2023) ‘Effects of Syllable Rate on Neuro-Behavioral Synchronization Across Modalities: Brain Oscillations and Speech Productions’, Neurobiology of Language, 4(2), pp. 344–360. Available at: 10.1162/nol_a_00102.

Herrmann, B., Buckland, C. and Johnsrude, I.S. (2019) ‘Neural signatures of temporal regularity processing in sounds differ between younger and older adults’, Neurobiology of Aging, 83, pp. 73–85. Available at: 10.1016/j.neurobiolaging.2019.08.028.

Herrmann, B. and Johnsrude, I.S. (2018) ‘Neural signatures of the processing of temporal patterns in sound’, Journal of Neuroscience, 38(24), pp. 5466–5477. Available at: 10.1523/JNEUROSCI.0346-18.2018.

Jaques, N., Gu, S., Bahdanau, D., Hernández-Lobato, J.M., Turner, R.E. and Eck, D. (2017) ‘Sequence Tutor: Conservative Fine-Tuning of Sequence Generation Models with KL-control’, in D. Precup and Y.W. Teh (eds) Proceedings of the 34th International Conference on Machine Learning. PMLR (Proceedings of Machine Learning Research), pp. 1645–1654. Available at: https://proceedings.mlr.press/v70/jaques17a.html (Accessed: 29 November 2022).

Keehn, B., Kadlaskar, G., McNally Keehn, R. and Francis, A.L. (2019) ‘Auditory Attentional Disengagement in Children with Autism Spectrum Disorder’, Journal of Autism and Developmental Disorders, 49(10), pp. 3999–4008. Available at: 10.1007/s10803-019-04111-z.

Kelly, S.P., Lalor, E.C., Reilly, R.B. and Foxe, J.J. (2006) ‘Increases in alpha oscillatory power reflect an active retinotopic mechanism for distracter suppression during sustained visuospatial attention’, Journal of Neurophysiology, 95(6), pp. 3844–3851. Available at: 10.1152/jn.01234.2005.

Kidd, G., Mason, C.R., Swaminathan, J., Roverud, E., Clayton, K.K. and Best, V. (2016) ‘Determining the energetic and informational components of speech-on-speech masking’, The Journal of the Acoustical Society of America, 140(1), pp. 132–144. Available at: 10.1121/1.4954748.

Kidd, G.Jr., Arbogast, T.L., Mason, C.R. and Walsh, M. (2002) ‘Informational Masking in Listeners with Sensorineural Hearing Loss’, Journal of the Association for Research in Otolaryngology, 3(2), pp. 107–119. Available at: 10.1007/s101620010095.

Kim, J. (2015) ‘Acoustic change complex: clinical implications’, Journal of Audiology & Otology, 19(3), pp. 120–124. Available at: 10.7874/jao.2015.19.3.120.

Kong, Y.-Y., Mullangi, A. and Ding, N. (2014) ‘Differential modulation of auditory responses to attended and unattended speech in different listening conditions’, Hearing research, 316, pp. 73–81.

Kraus, N., Thompson, E.C., Krizman, J., Cook, K., White-Schwoch, T. and LaBella, C.R. (2016) ‘Auditory biological marker of concussion in children’, Scientific Reports, 6(1), pp. 1–10. Available at: 10.1038/srep39009.

Krizman, J. and Kraus, N. (2019) ‘Analyzing the FFR: a tutorial for decoding the richness of auditory function’, Hearing Research, 382. Available at: 10.1016/j.heares.2019.107779.

Kuwada, S., Anderson, J.S., Batra, R., Fitzpatrick, D.C., Teissier, N. and D’Angelo, W.R. (2002) ‘Sources of the scalp-recorded amplitude-modulation following response’, Journal of the American Academy of Audiology, 13(4), pp. 188–204. Available at: 10.1055/S-0040-1715963.

de Larrea-Mancera, E.S.L., Philipp, M.A., Stavropoulos, T., Carrillo, A.A., Cheung, S., Koerner, T.K., Molis, M.R., Gallun, F.J. and Seitz, A.R. (2022) ‘Training with an auditory perceptual learning game transfers to speech in competition’, Journal of Cognitive Enhancement, 6(1), pp. 47–66. Available at: 10.1007/s41465-021-00224-5.

McKenna, L., Handscomb, L., Hoare, D.J. and Hall, D.A. (2014) ‘A scientific cognitive-behavioral model of tinnitus: Novel conceptualizations of tinnitus distress’, Frontiers in Neurology, 5(OCT). Available at: 10.3389/FNEUR.2014.00196/FULL.

Müller, N., Schlee, W., Hartmann, T., Lorenz, I. and Weisz, N. (2009) ‘Top-down modulation of the auditory steady-state response in a task-switch paradigm’, Frontiers in Human Neuroscience, 3(FEB). Available at: 10.3389/NEURO.09.001.2009/FULL.

Nolan, H., Whelan, R. and Reilly, R.B. (2010) ‘FASTER: Fully Automated Statistical Thresholding for EEG artifact Rejection’, Journal of neuroscience methods, 192(1), pp. 152–162. Available at: 10.1016/J.JNEUMETH.2010.07.015.

O’Connor, K. (2012) ‘Auditory processing in autism spectrum disorder: a review’, Neuroscience & Biobehavioral Reviews, 36(2), pp. 836–854. Available at: https://www.sciencedirect.com/science/article/pii/S0149763411002065 (Accessed: 18 March 2025).

Parthasarathy, A., Hancock, K.E., Bennett, K., Degruttola, V. and Polley, D.B. (2020) ‘Bottom-up and top-down neural signatures of disordered multi-talker speech perception in adults with normal hearing’, eLife, (9), p. e51419. Available at: 10.7554/eLife.51419.

Picton, T.W., John, M.S., Dimitrijevic, A. and Purcell, D. (2003) ‘Human auditory steady-state responses’, International Journal of Audiology, 42(4), pp. 177–219. Available at: 10.3109/14992020309101316.

Rosen, S. (1992) ‘Temporal information in speech: acoustic, auditory and linguistic aspects.’, Philosophical transactions of the Royal Society of London. Series B, Biological sciences. Philos Trans R Soc Lond B Biol Sci, pp. 367–373. Available at: 10.1098/rstb.1992.0070.

Schüller, A., Schilling, A., Krauss, P., Rampp, S. and Reichenbach, T. (2023) ‘Attentional modulation of the cortical contribution to the frequency-following response evoked by continuous speech’, Journal of Neuroscience, 43(44), pp. 7429–7440. Available at: 10.1523/JNEUROSCI.1247-23.2023.

Shinn-Cunningham, B., amplification, V.B.-T. in and 2008, undefined (2008) ‘Selective attention in normal and impaired hearing’, journals.sagepub.com, 12(4), pp. 283–299. Available at: 10.1177/1084713808325306.

Shinn-Cunningham, B., Varghese, L., Wang, L. and Bharadwaj, H. (2017) ‘Individual Differences in Temporal Perception and Their Implications for Everyday Listening’, pp. 159–192. Available at: 10.1007/978-3-319-47944-6_7.

Shinn-Cunningham, Barbara, Varghese, Leonard, Wang, Le, Bharadwaj, Hari, Shinn-Cunningham, B, Varghese, Á.L., Wang, Á.L., Varghese, L, Wang, L and Bharadwaj, H (2017) ‘Individual differences in temporal perception and their implications for everyday listening’, Springer, p. 61. Available at: 10.1007/978-3-319-47944-6_7.

Sollini, J., Poole, K.C., Blauth-Muszkowski, D. and Bizley, J.K. (2022) ‘The role of temporal coherence and temporal predictability in the build-up of auditory grouping’, Scientific Reports 2022 12:1, 12(1), pp. 1–10. Available at: 10.1038/s41598-022-18583-0.

Southwell, R., Baumann, A., Gal, C., Barascud, N., Friston, K. and Chait, M. (2017) ‘Is predictability salient? A study of attentional capture by auditory patterns’, Philosophical Transactions of the Royal Society B: Biological Sciences, 372(1714). Available at: 10.1098/RSTB.2016.0105.

Stevens, K.N. (1971) ‘Sources of Inter- and Intra-Speaker Variability in the Acoustic Properties of Speech Sounds’, in A. Rigault and R. Charbonneau (eds) Proceedings of the seventh International Congress of Phonetic Sciences. De Gruyter, pp. 206–232. Available at: 10.1515/9783110814750-014.

Studebaker, G.A. (1985) ‘A “rationalized” arcsine transform.’, Journal of Speech and Hearing Research, 28(3), pp. 455–462. Available at: 10.1044/JSHR.2803.455.

Swaminathan, J., Mason, C.R., Streeter, T.M., Best, V., Roverud, E. and Kidd, G. (2016) ‘Role of Binaural Temporal Fine Structure and Envelope Cues in Cocktail-Party Listening’, The Journal of Neuroscience, 36(31), p. 8250. Available at: 10.1523/JNEUROSCI.4421-15.2016.

Tierney, A., Rosen, S., Psychology, F.D.-J. of E. and 2020, undefined (2020) ‘Speech-in-speech perception, nonverbal selective attention, and musical training.’, psycnet.apa.org, 46(5), pp. 968–979. Available at: 10.1037/xlm0000767.

Tremblay, K.L., Pinto, A., Fischer, M.E., Klein, B.E.K., Klein, R., Levy, S., Tweed, T.S. and Cruickshanks, K.J. (2015) ‘Self-Reported Hearing Difficulties among Adults with Normal Audiograms: The Beaver Dam Offspring Study’, Ear and Hearing, 36(6), pp. e290–e299. Available at: 10.1097/AUD.0000000000000195.

Undurraga, J.A., Haywood, N.R., Marquardt, T. and McAlpine, D. (2016) ‘Neural Representation of Interaural Time Differences in Humans—an Objective Measure that Matches Behavioural Performance’, Journal of the Association for Research in Otolaryngology, 17(6), pp. 591–607. Available at: 10.1007/s10162-016-0584-6.

Varghese, L., Bharadwaj, H.M. and Shinn-Cunningham, B.G. (2015) ‘Evidence against attentional state modulating scalp-recorded auditory brainstem steady-state responses’, Brain Research, 1626, pp. 146–164. Available at: 10.1016/j.brainres.2015.06.038.

Vercammen, C., Goossens, T., Undurraga, J., Wouters, J. and Van Wieringen, A. (2018) ‘Electrophysiological and behavioral evidence of reduced binaural temporal processing in the aging and hearing impaired human auditory system’, Trends in Hearing, 22. Available at: 10.1177/2331216518785733.

Verhulst, S., Altoè, A. and Vasilkov, V. (2018) ‘Computational modeling of the human auditory periphery: Auditory-nerve responses, evoked potentials and hearing loss’, Hearing Research, pp. 55–75. Available at: 10.1016/j.heares.2017.12.018.

Wang, L., Bharadwaj, H. and Shinn-Cunningham, B. (2019) ‘Assessing Cochlear-Place Specific Temporal Coding Using Multi-Band Complex Tones to Measure Envelope-Following Responses’, Neuroscience, 407, pp. 67–74. Available at: 10.1016/j.neuroscience.2019.02.003.

Whitton, J.P., Hancock, K.E., Shannon, J.M. and Polley, D.B. (2017) ‘Audiomotor Perceptual Training Enhances Speech Intelligibility in Background Noise’, Current Biology, 27(21), pp. 3237–3247.e6. Available at: 10.1016/j.cub.2017.09.014.

Wöstmann, M., Herrmann, B., Maess, B. and Obleser, J. (2016) ‘Spatiotemporal dynamics of auditory attention synchronize with speech’, Proceedings of the National Academy of Sciences of the United States of America, 113(14), pp. 3873–3878. Available at: 10.1073/pnas.1523357113.

Zhu, L., Bharadwaj, H., Xia, J. and Shinn-Cunningham, B. (2013) ‘A comparison of spectral magnitude and phase-locking value analyses of the frequency-following response to complex tones’, The Journal of the Acoustical Society of America, 134(1), pp. 384–395. Available at: 10.1121/1.4807498.

