## Supplemental Audio Files for "The Slowest Timescales of Neural Synchronization Reveal the Strongest Influence of Auditory Distraction": Supplementary_Material.docx

### 1 Supplementary Data

Exemplar audio files are included as supplementary data. Descriptions of the included files:

Target_Random-Pattern.wav An example target random-pattern stimulus used in the EEG sessions, corresponding to the illustration in Figure 1C.

Target_Random-Random.wav An example target random-random stimulus used in the EEG sessions.

Target_Pattern.wav An example target pattern stimulus used in the at-home testing, corresponding to the illustration in Figure 2B.

Melodic_Distractor_1.wav An example melodic distractor stimulus corresponding to the top panel in Figure 2A and in Figure 2B

Melodic_Distractor_2.wav An example melodic distractor stimulus corresponding to the middle panel in Figure 2A

Melodic_Distractor_3.wav An example melodic distractor stimulus corresponding to the bottom panel in Figure 2A

MatchedNoise_Distractor_1.wav An example noise distractor stimulus corresponding to the illustration in Figure 4F, and matched to Melodic_Distractor_1.wav

MatchedNoise_Distractor_2.wav An example noise distractor stimulus matched to Melodic_Distractor_2.wav

MatchedNoise_Distractor_3.wav An example noise distractor stimulus matched to Melodic_Distractor_3.wav
